# Proton tunneling at the ryanodine receptor Ca^2+^ activation site provides temperature-invariant noise for robust Ca^2+^-induced Ca^2+^ release

**DOI:** 10.64898/2026.04.08.717335

**Authors:** Alexander V. Maltsev, Edward G. Lakatta, Victor A. Maltsev

## Abstract

Ryanodine receptor (RyR)-mediated Ca^2+^-induced Ca^2+^ release (CICR) is a regenerative trigger-to-release mechanism used in diverse cell types. RyRs amplify small Ca^2+^ signals into large, localized surges that drive autonomous oscillation, excitation–contraction coupling, pulsatile secretion, neurotransmitter release, or memory formation. Robust RyR function depends on the probability of RyR recruitment remaining within an operating range as physiological conditions change. This raises a basic question: what stabilizes the microscopic Ca^2+^-sensitive opening step of RyR across temperature and cellular state? Here we tested that proton tunneling at the conserved RyR2 Ca^2+^-activation site contributes a temperature-stable stochastic component to channel opening. To this end we developed a multiscale quantum-structural pipeline that estimates the per-channel modulation amplitude from cryo-EM geometry under stated assumptions. We applied this pipeline to a rabbit sinoatrial node cell model with explicit Ca^2+^ release unit architecture. Holding this RyR quantum noise constant across 25–37 ^∘^C preserves pacemaker rhythmicity at low temperature, whereas a *Q*_10_-scaled classical noise results in irregular pacemaking activity as amplitude falls below the coherence-resonance range. Experimentally in isolated rabbit sinoatrial node cells, Ca^2+^ transient rate increased with temperature, but inter-transient interval variability did not, consistent with our numerical model predictions. These results support a model in which proton tunneling stabilizes the regenerative RyR-mediated CICR step. In autonomous oscillators, such as sinoatrial node cells, this manifests as robust rhythmicity. In triggered systems, it is expected to stabilize CICR gain, release synchrony, and release fidelity. The importance of this shared microscopic mechanism is evidenced by the conservation of the Ca^2+^-activation site across RyR isoforms for >600 million years of evolution.

**Significance Statement:** This work suggests that RyR-based CICR networks can harness quantum-conditioned stochasticity. Proton tunneling at a conserved RyR activation site contributes a temperature-stable source of microscopic variability, and CICR amplifies that variability into robust physiological output. In pacemaker cells, this appears as stable rhythmic timing. More broadly, it provides a concrete route by which atomic-scale quantum events can influence macroscopic cellular function.

## Introduction

Ryanodine receptor (RyR)-mediated Ca^2+^-induced Ca^2+^ release (CICR) is a regenerative trigger-to-release mechanism used in diverse physiological settings. CICR helps organize rhythmic firing in the sinoatrial node (SAN) [1,2], amplifies L-type Ca^2+^ influx into cell-wide sarcoplasmic reticulum release in ventricular myocytes [3,4,5], and contributes to pulsatile insulin secretion in pancreatic *β*-cells [6,7], pacemaker activity in interstitial cells of Cajal [8,9], and somatodendritic signaling in cerebellar Purkinje neurons [10,11]. The physiological output differs across these systems, but robust function depends on the Ca^2+^-sensitive RyR recruitment step remaining within an optimal operating range. Because regenerative release operates near an ignition threshold, even modest shifts in the microscopic opening step can have macroscopic consequences.

A local stochastic opening can recruit neighboring channels and be amplified into a Ca^2+^ spark [3,12,5] or larger release event [13,14]. In SAN cells, this regenerative architecture makes rhythmicity sensitive to stochastic fluctuations [15,16], and both stochastic resonance and coherence resonance predict that noise can improve regularity only within a limited amplitude window [17,18,19]. Thus, stochasticity is not a harmful disturbance, but an essential component of physiological function [14]. This raises a basic question: what stabilizes the Ca^2+^-sensitive RyR opening step as temperature and cellular state change?

Lindner and colleagues framed this as a foundational challenge. The origins of noise in excitable systems are as diverse as the physical basis of excitability itself, and identifying them requires committing to a specific physical substrate [18]. A recent review of cardiac noise echoes the same point in the SAN context, identifying the physical source of SR Ca^2+^ release noise as an explicit open question [14], and analogous noise-calibration questions remain open across fields as far apart as inner ear mechanotransduction, vertebrate segmentation, and plant guard cell signaling.

A longstanding biophysical anomaly suggests that the opening step of RyR may be governed by different physics than its closing transitions. In single-channel recordings of cardiac RyR across 5–23 ^∘^C, the closing rate showed the strong temperature dependence expected for classical protein conformational changes, whereas the opening rate was nearly temperature-independent [20]. The electron-conformational model of RyR [21,22] proposes the initial activation as a transition between two electronic states (the “pseudospin” jump). This transition was mathematically treated as a quantum tunneling event between two potential wells. If this limiting step indeed occurs by tunneling, the opening transition would depend primarily on donor-acceptor geometry and tunneling particle (proton or electron) mass rather than on thermal energy.

This possibility has a direct systems-level implication. A classical stochastic input should drift with temperature, metabolic state, and other perturbations, whereas a tunneling-linked component of RyR opening could remain comparatively stable. Stabilization of this microscopic ignition step could preserve the probability of spark recruitment even as other cellular processes scale classically. This would produce more stable beat-to-beat timing in autonomous oscillators such as the SAN and more robust excitation-contraction coupling gain, release latency, and release synchrony in triggered systems such as ventricular myocytes. As used here, “quantum noise” does not imply long-lived coherence at the cellular scale but rather a stochastic modulation of RyR opening probability arising from tunneling at the Ca^2+^-activation interface. This shifts the role of quantum biology in this context from its traditional emphasis on reaction speed in enzyme catalysis and electron transport [23,24,25,26] to the control of biological variability in a nonlinear excitable system.

The SAN pacemaker offers a particularly informative test of this mechanism. The SAN, as an autonomous oscillator, reports cycle-to-cycle variability, reflecting the amplitude of the organizing noise rather than the AP firing rate [27,17,18,28]. If RyR opening carries a temperature-stable stochastic component (Fig. 1A), then temperature-dependent membrane transport, SERCA activity, and diffusion should still shift the firing interval, but the variability of interbeat timing should change much less. The SAN readout therefore reports on the stability of the regenerative Ca^2+^ release step that organizes rhythm.

**Figure 1.**
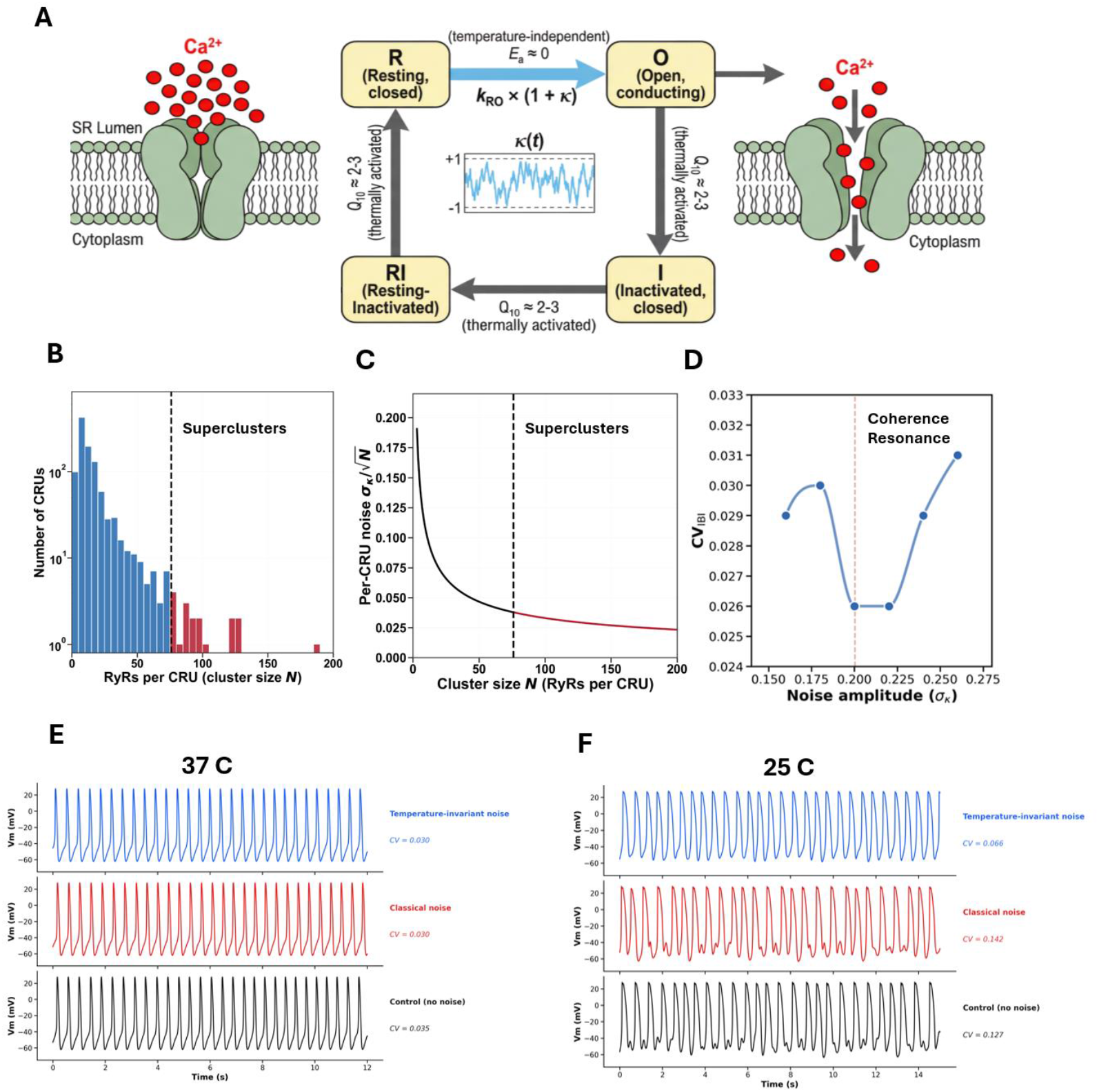
Temperature-independent noise maintains rhythmic regularity in a SAN cell model. (**A**) RyR2 four-state gating scheme (R, O, I, RI). Ca^2+^-activated opening (R→O) is temperature-independent (*E*_*a*_ ≈ 0) and modulated by a stochastic factor κ(*t*); other transitions are thermally activated (*Q*_10_ ≈ 2–3). (**B**) Distribution of CRU sizes (RyRs per cluster, *N*) from super-resolution imaging; dashed line marks the threshold separating typical clusters from large superclusters (red). (**C**) Per-CRU noise amplitude 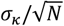 versus *N*, showing that larger superclusters experience smaller effective per-CRU noise. (**D**) Coherence-resonance dose-response at 37 °C: CV_IBI_ versus *σ*_*κ*_ traces a U-shape with a minimum near *σ*_*κ*_ ≈ 0.20 (dashed line; SI Table S6). (**E**) Simulated membrane potential at 37 °C for no-noise control (top), *Q*_10_-scaled classical noise (*σ* = 0.20; middle), and temperature-invariant “quantum” noise (*σ* = 0.20; bottom). All three are rhythmic. (**F**) Same conditions at 25 °C: control and classical noise become arrhythmic, whereas temperature-invariant noise remains rhythmic, consistent with the classical amplitude falling below the coherence-resonance window at low temperature.

Cryo-EM structures of RyR1 and RyR2 identify a conserved Ca^2+^-activation site formed by E3848, E3922, and T4931 [29,30,31]. These residues are conserved across mammalian RyR isoforms and across deep animal evolution [4,31], consistent with a shared microscopic mechanism whose physiological importance depends on release architecture. Here we report evidence that proton tunneling at this conserved site contributes a temperature-stable stochastic component to RyR opening, and that this is important for physiological function, i.e. robust rhythmicity of cardiac pacemaker cells. To test this, our study combined structural analysis, quantum modeling, single-cell simulations, and Ca^2+^ imaging in isolated rabbit SAN cells.

## Results

### the coherence resonance analysis above

At 37 ^∘^C, the single-cell SAN model [32,33] (Methods) with four-state RyR kinetics and super-clustering (Fig. 1A,B) generates rhythmic APs. However, when *Q*_10_ temperature scaling is applied to all classically temperature-dependent processes (Methods), the model exhibits irregular AP firing at 25 ^∘^C with high CV of inter-transient intervals (CV_IBI_; Fig. 1F; SI Appendix, Table S1). An additional noise term was added to the RyR2 Ca^2+^-dependent opening rate *k*_oCa_ (Fig. 1C, Methods, Eq. 3), and its amplitude *σ*_*κ*_ was scanned at 37 ^∘^C. The result is U-shaped: at low amplitudes CV_IBI_ is high, at an intermediate amplitude it reaches a minimum, and at high amplitudes it rises again (Fig. 1D; SI Appendix, Table S6). In a separate set of temperature-sweep simulations, two versions of this noise were compared at 25 ^∘^C and 37 ^∘^C: temperature-independent noise (*σ*_*κ*_ = 0.20, constant, labeled “quantum”) and *Q*_10_-scaled noise (*σ*_*κ*_ = 0.20 at 37 ^∘^C declining to 0.061 at 25 ^∘^C, labeled “classical”), alongside a no-noise control (*κ* = 0). At 37 ^∘^C all three conditions produce comparable CV_IBI_ (0.030, 0.030, and 0.035 for quantum, classical, and control respectively). At 25 ^∘^C they diverge: quantum noise maintains low CV_IBI_ (0.066), while classical (0.142) and control (0.127) show substantially higher variability (Fig. 1E,F). Only the temperature-invariant noise condition maintained low CV_IBI_ at both temperatures.

### Experimental Ca^2+^ transients show temperature-invariant variability

Ca^2+^-dependent fluorescence recordings (Fig. 2A,B) from 17 isolated rabbit SAN cells, each recorded at both 25 ^∘^C and 37 ^∘^C (paired design; 100 Hz frame rate; Methods), were analyzed. All 17 cells fired rhythmically at both temperatures. Beat rate increased from 0.96 ± 0.13 Hz at 25 ^∘^C to 1.60 ± 0.18 Hz at 37 ^∘^C (*p* = 1.17 × 10^−10^, paired t-test, *n* = 17). In contrast, CV_IBI_ did not change with temperature: 0.096 ± 0.054 at 25 ^∘^C vs. 0.067 ± 0.035 at 37 ^∘^C (*p* = 0.050, paired t-test, *n* = 17; Fig. 2C–F). Beat rate scaled with temperature while CV_IBI_ did not.

**Figure 2.**
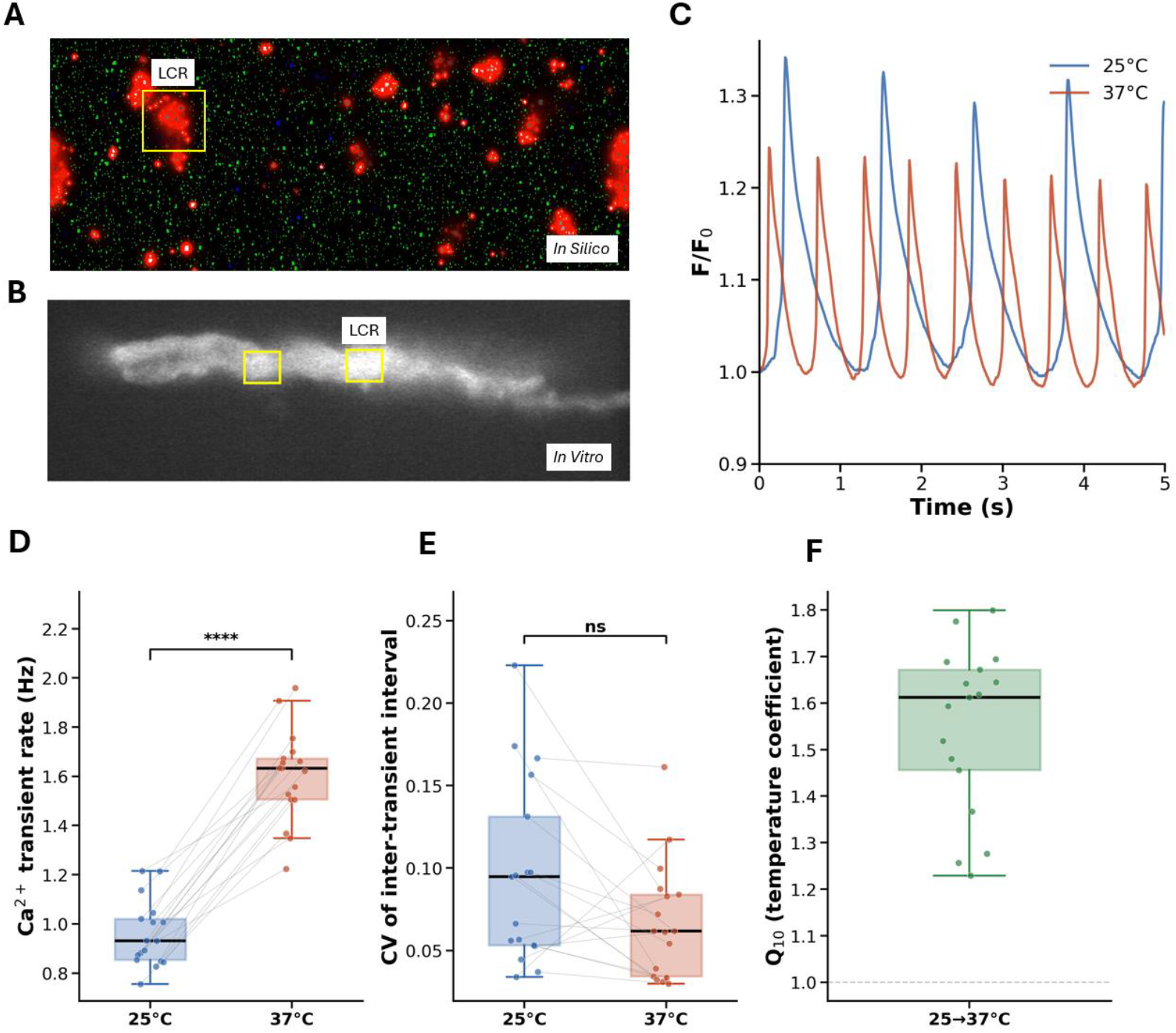
Model and experimental Ca^2+^ transients show temperature-invariant variability. (**A**) Representative *in silico* image of local Ca^2+^ release from the SAN model, with an LCR event highlighted (yellow box). (**B**) Representative *in vitro* Fluo-4 fluorescence image of an isolated rabbit SAN cell with LCR regions highlighted. (**C**) Whole-cell Ca^2+^ transients (*F*/*F*_0_) from a single cell at 25 °C (blue) and 37 °C (red). (**D**) Beat rate at 25 °C versus 37 °C (*n* = 17, paired) increased significantly with warming (paired t-test, *p* = 1.17 × 10^−10^). (**E**) CV of inter-transient intervals for the same cells did not differ between temperatures (ns; *p* = 0.050). (**F**) Cell-by-cell *Q*_10_ for beat rate (25 °C → 37 °C), mean 1.55 ± 0.17 (median 1.61). Box plots show median and IQR; thin lines connect paired measurements.

### Cryo-EM structures reveal short hydrogen-bond contacts at the activation site

The O⋯O distances at the Ca^2+^ activation site were measured across 12 recent open-access cryo-EM structures spanning human and porcine RyR2 in open and closed states, plus rabbit RyR1 as a cross-isoform check (SI Appendix, Table S3). The open and closed states differ systematically. In the open state (Ca^2+^ bound), six structures show E3848–E3922 O⋯O distances in H-bonding range (2.88–3.82 Å, mean 3.31 ± 0.35 Å), while three show non-H-bonding distances (4.58–6.19 Å; Fig. 3A,B). In the closed state (Ca^2+^ absent), all distances exceed 5 Å. Short carboxylate–carboxylate contacts appear only when Ca^2+^ is bound and the channel is open. Two structures approach 3 Å: 6JIY at 2.88 Å and 5TAL (RyR1) at 3.01 Å. For comparison, neutron crystallography has confirmed proton sharing between carboxylates at O⋯O distances of 2.43–2.55 Å [34,35,36,37]. The cryo-EM distances are 0.33–0.45 Å longer than the confirmed tunneling range. B-factor analysis confirms that the activation site is consistently more ordered than the protein average (*B*_site_/*B*_global_ = 0.57 ± 0.16; *p* < 10^−3^, Wilcoxon signed-rank test against 1, n = 12; Fig. 3C; SI Appendix, Table S3). The single outlier (7U9R, ratio ≈ 1.0) is the PKA-phosphorylated open-state structure in which the E3848–E3922 distance expands to 6.19 Å and Ca^2+^ no longer bridges the two carboxylates, consistent with Ca^2+^-mediated rigidity at the activation site. Two pK_*a*_ predictors bracket E3922 protonation at 3–65% at pH 7.0 (SI Appendix, Tables S4–S5).

**Figure 3.**
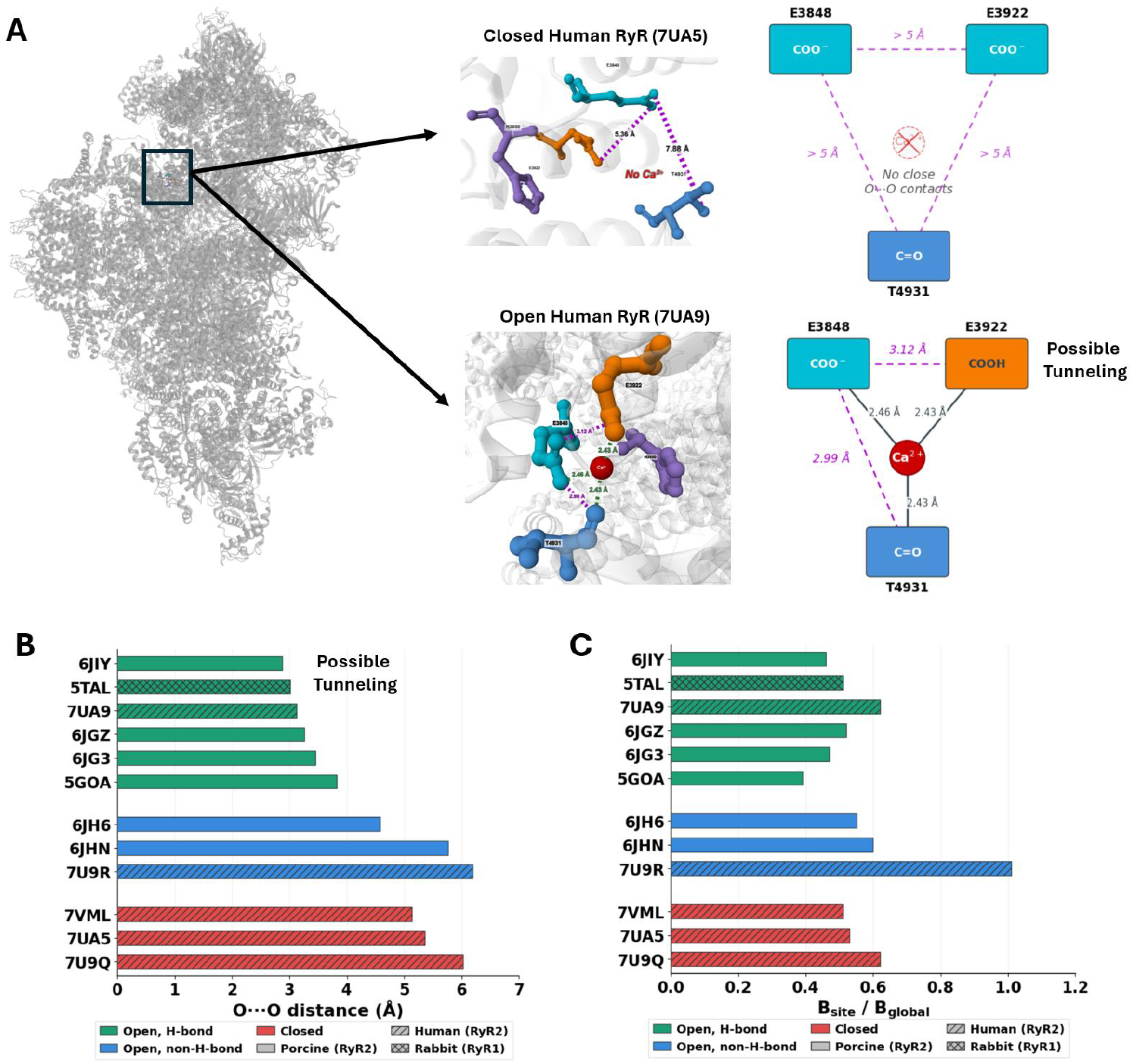
Cryo-EM structures reveal short hydrogen-bond contacts at the RyR Ca^2+^-activation site only in the open, Ca^2+^-bound state. (**A**) Overall RyR architecture with the activation site boxed. Closed human RyR2 (7UA5; top) shows all O ⋯ O distances > 5 Å with no bound Ca^2+^. Open human RyR2 (7UA9; bottom) shows Ca^2+^ coordinating both carboxylates and T4931, with the E3848–E3922 distance contracting to 3.12 Å—geometry compatible with a shared proton and possible tunneling. Schematics summarize the donor-acceptor geometry. (**B**) E3848–E3922 O ⋯ O distances across 12 cryo-EM structures (SI Table S3): open H-bonding (green; 2.88–3.82 Å), open non-H-bonding (blue; 4.58–6.19 Å), and closed (red; > 5 Å). Hatching denotes species/isoform. (**C**) Ratio of activation-site mean B-factor to protein-wide mean B-factor (*B*_si*t*e_/*B*_global_) for the 12 structures in panel B. Values below 1 indicate the site is more ordered than the protein average (mean 0.57 ± 0.16; *p* < 0.001, Wilcoxon signed-rank test against 1, *n* = 12). The sole outlier 7U9R (≈ 1.0) is the PKA-phosphorylated structure in which Ca^2+^ no longer bridges E3848–E3922, consistent with Ca^2+^-mediated rigidity at the site.

### Proton tunneling at the activation site produces temperature-invariant noise within the coherence resonance window

The cryo-EM O⋯O distances are longer than the range where tunneling has been directly confirmed. A four-step quantum-mechanical pipeline was used to test whether thermal fluctuations can compress the activation site into tunneling-ready configurations often enough to produce noise within the coherence resonance window (Fig. 4A,B). The O⋯O distance fluctuates thermally around its equilibrium value. Ca^2+^-coordination-weighted Boltzmann rotamer sampling across all six open H-bonding structures yields 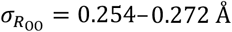 (mean 0.264 ± 0.006 Å), bounded from below by the Herschlag thermodynamic limit (≥ 0.06 Å; [38]). These fluctuations occasionally compress the O⋯O distance into the tunneling-competent regime (∼ 2.50 Å), where the tunneling splitting *ΔE* rises by orders of magnitude (Fig. 4B,C), following the “tunneling-ready configuration” approach used in enzyme catalysis [23]. A saturating transfer function converts these intermittent splitting spikes into a bounded noise signal κ(*t*), and two-tier coarse-graining [39] over the 4 ms simulation update window yields *σ*_κ,eff_ ≈ 0.20 (Fig. 4D,E; SI Appendix, Methods and Table S6); this value is robust across a reasonable range of slow-tier parameters (Fig. 4D). This amplitude falls within the coherence resonance window identified in the coherence resonance analysis above. The amplitude changes by <1% across 5–42 ^∘^C, as the saturating transfer function absorbs the modest 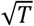 scaling of 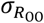 (Fig. 4F; SI Appendix, Table S6).

**Figure 4.**
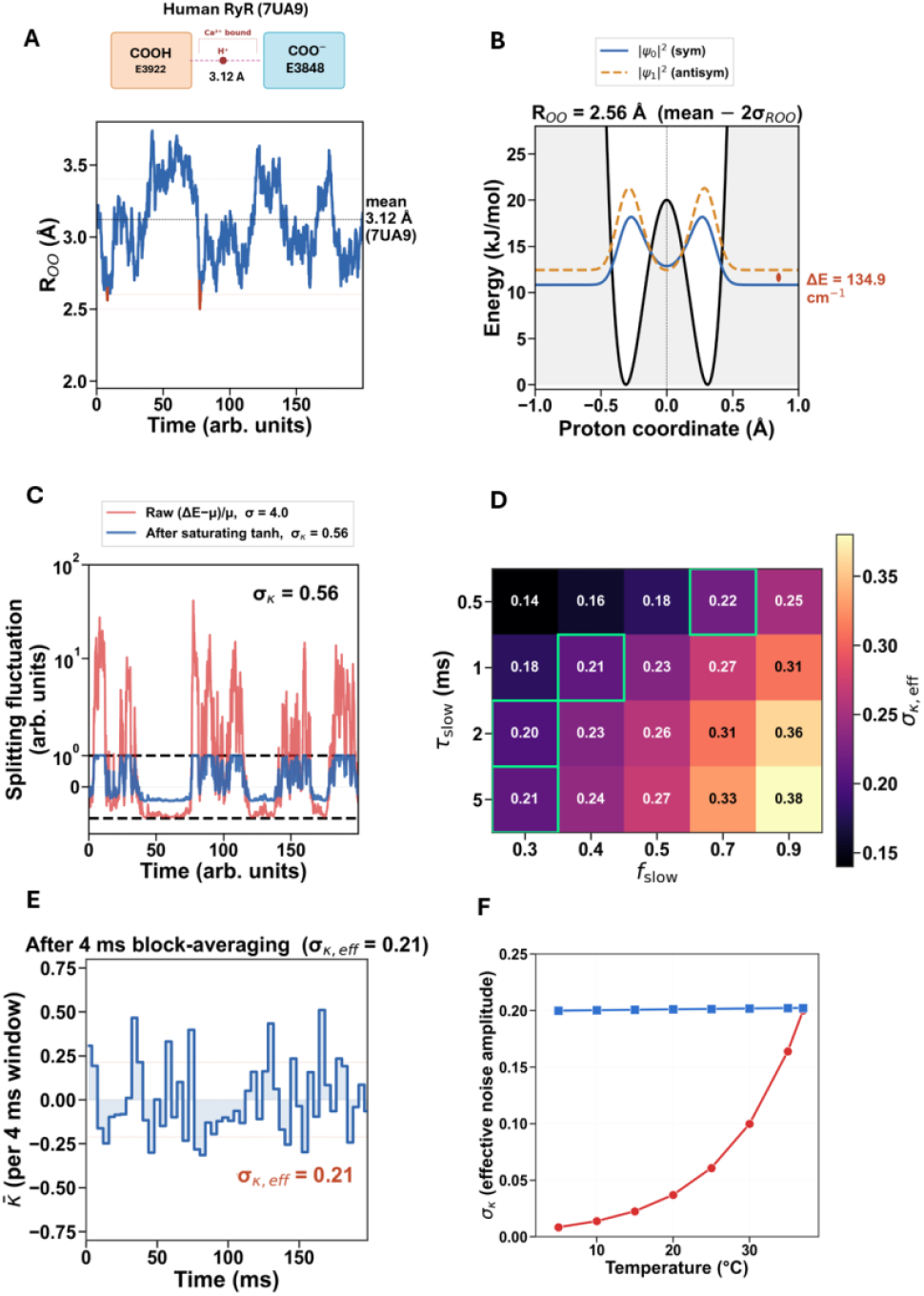
Quantum pipeline converts cryo-EM geometry into a temperature-invariant per-channel noise amplitude. (**A**) Top: schematic of the E3922 (COOH) – E3848 (COO^−^) pair in open human RyR2 (7UA9), mean *R*_*00*_ = 3.12 Å. Bottom: representative two-tier Ornstein–Uhlenbeck trajectory of *R*_*00*_; excursions to short distances (red) transiently reach the tunneling-competent regime. (**B**) Solution of the 1D Schrödinger equation (Eq. 1) for a proton in a quartic double-well at *R*_*00*_ = 2.56 Å. Symmetric |*ψ*_0_|^2^ (blue) and first-excited-state |*ψ*_1_|^2^ (orange) are separated by *ΔE* = 134.9 cm^−1^. (**C**) Raw splitting fluctuation (red, *σ* = 4.0) and its bounded form after the saturating tanh transfer function (blue, *σ*_*κ*_ = 0.56). (**D**) Sensitivity of *σ*_κ,eff_ to OU parameters *τ*_slow_ (0.5–5 ms) and *f*_slow_ (0.3–0.9). Green-outlined cells fall within the coherence-resonance window; high-*τ*_slow_ / high-*f*_slow_ combinations are disfavored by Ca^2+^ coordination at the site. (**E**) Block-averaged modulation 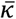 over the 4 ms update window, yielding *σ*_κ,eff_ = 0.20 at reference parameters (*τ*_slow_ = 1 ms, *f*_slow_ = 0.4). (**F**) *σ*_*κ*_ versus temperature for quantum (blue) and *Q*_10_-scaled classical (red) noise. The quantum amplitude varies < 1% across 5–42 °C because the saturating transfer function absorbs the modest 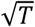 scaling of 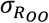, whereas the classical amplitude falls steeply with cooling.

## Discussion

Our results point to a mechanism that can stabilize the ignition step of RyR-mediated Ca^2+^ release, where proton tunneling at the conserved RyR Ca^2+^ activation site adds a temperature-stable stochastic component to channel opening. We used an example of SAN pacemaker cell function to illustrate that such tunneling can ensure robust temperature-invariant regular beating. Cryo-EM structures identify donor-acceptor geometry at the open RyR activation site that is compatible with proton transfer. The quantum pipeline converts that geometry into a per-channel noise amplitude. In the SAN cell model, holding this RyR-side noise constant preserves rhythmic regularity at low temperature, while a *Q*_10_-scaled classical noise term falls below the effective range and regularity degrades. Recordings from isolated rabbit SAN cells show the same separation at the cell level. Mean Ca^2+^ transient rate increases from 25 ^∘^C to 37 ^∘^C, while interval variability remains stable. Together these findings support the idea that the RyR opening step carries a distinct temperature-invariant stochastic component that stabilizes regenerative CICR.

The physiological importance of the mechanism follows from where it acts. RyR clusters sit at the transition between local and self-amplifying release [5,13,14]. Small changes in RyR Ca^2+^ sensitivity can therefore determine whether a local event dies out or becomes a spark and with further amplification become a larger event such as a local Ca^2+^ release (LCR; multi-spark event) [5,40]. In SAN cells, action potential timing depends on early diastolic release events from the CRUs that engage the membrane clock [27,28,32,41]. Stabilizing the RyR opening step stabilizes the rhythm. This provides a microscopic origin for the LCR timing dispersion that Monfredi et al. linked to intrinsic cycle-length variability in isolated SAN cells [15]. AP firing rate still changes with temperature because membrane currents, SERCA activity, diffusion, SR loading, and other parts of the pacemaker system continue to scale classically.

This principle extends beyond the SAN. The E3848–E3922–T4931 Ca^2+^ activation triad is conserved across RyR isoforms and across 600 million years of evolution [4,31], pointing to a shared microscopic mechanism at the Ca^2+^myocytes [5], and reliable pulsatile or rhythmic Ca^2+^ recruitment in pancreatic *β*-cells, interstitial cells of Cajal, and Purkinje neurons [6,7,8,9,10,11].

RyR is a particularly effective site for this mechanism because CICR amplifies microscopic fluctuations into macroscopic events. Heterogeneity in CRU size sharpens that effect because a limited subset of larger superclusters can exert a disproportionate influence on cell-level behavior [32]. Unlike the decoherence-compatible tunneling studied in enzyme catalysis and DNA proton transfer [23,42,43], the functional output here is not a classical reaction rate but a stochastic modulation: atomic-scale variability at the activation site shapes variability-dependent whole-cell physiology.

Conceptually, this places RyR-based CICR in a class of biological architectures that act as quantum-to-classical transducers. The microscopic, quantum-conditioned transition probability is amplified through a regenerative threshold element into a macroscopic outcome. The quantum step does not encode the heartbeat, but instead it perturbs the statistics of an opening transition that the cell is already poised to amplify, and the cell’s regenerative architecture does the rest. The same physics is well established outside biology. In mesoscopic conductors, the noise of any tunnel junction is set by barrier geometry and is independent of temperature [44]; in scanning tunneling microscopy of single iron atoms, quantum tunneling generates a stochastic resonance signal whose amplitude does not vanish as temperature approaches zero [45]. Enzymology has documented for four decades that proteins can hold donor–acceptor geometry rigid enough to render hydrogen tunneling rates temperature-independent [23]. We propose that the conserved Ca^2+^-activation site of RyR has evolved to do the same thing for a different purpose, i.e. not to accelerate a catalytic step, but to inject a temperature-stable stochastic input into an excitable amplifier. This is consistent with the broader principle, articulated by Noble and others, that organisms harness rather than merely tolerate stochasticity [46]. Our results extend this and suggest that conserved molecular geometries may be selected not only for their mean kinetics but for the statistics of fluctuation they deliver to higher-level control loops.

The same design problem appears in several systems with an unexplained noise-calibration requirement. Inner ear hair bundles operate at a Hopf bifurcation whose tuning requires noise amplitude just below thermal [47]. Vertebrate segmentation clock cells require a residual intrinsic noise floor for reproducible somite formation [48]. Plant guard cells encode stomatal aperture in Ca^2+^ oscillations within a defined frequency [49]. We propose that these are instances of a single principle: a regenerative or threshold-crossing dynamics requires a noise reference point that holds steady against temperature and metabolic state, and biology has few physical substrates that can supply one. Geometry-set tunneling at a conserved hydrogen-bond site is one such substrate, and the present work offers it as a worked example.

Several long-standing anomalies in cardiac physiology are compatible under this framework. Hibernating ground squirrels maintain slow but regular cardiac rhythm at body temperatures of 2–5 ^∘^C with both autonomic limbs withdrawn [50]. Every classical pacemaker mechanism (HCN currents, L-type Ca^2+^ current, SERCA, NCX) shows *Q*_10_autonomic input. A temperature-stable noise floor sustained by molecular geometry rather than by thermally activated rates is the kind of mechanism that could sustain it. A zebrafish study of two-clock pacemaking found that the Ca^2+^ clock contribution is independent of temperature while the membrane clock is not [51], an asymmetry consistent with the same interpretation of temperature-invariant noise source built into the molecular geometry of RyR. Crutchfield et al. recently used stochastic thermodynamics to show formally that voltage-gated ion channels operating under realistic conditions violate detailed balance, concluding that the underlying fluctuations cannot be accounted for by classical thermal noise alone [52].

Future studies follow directly from this framework. Pressure perturbations and targeted changes at the E3848–E3922–T4931 activation site should shift the tunneling geometry and move the system along the coherence resonance curve. QM/MM simulations with explicit Ca^2+^ and solvent can refine the proton transfer pathway and quantify the effective noise term at the native site. Deuterium substitution at the activation site should change the noise amplitude and directly affect the quantum tunneling rate. In ventricular myocytes, the model predicts relative stability of the RyR-dependent component of excitation-contraction coupling gain, spark recruitment, and release synchrony after controlling for trigger current and SR load. Other RyR- and *IP*_3_R-dependent excitable systems operating near coherence resonance should show a similar dissociation between temperature-dependent mean rate and temperature-stable interval variability (such as pancreatic *β*-cell insulin oscillations, interstitial cells of Cajal, Purkinje neuron pacemaking).

Thus, our data support a model in which proton tunneling stabilizes the regenerative RyR-mediated CICR step. In SAN cells that stabilization appears as robust rhythmicity. In other RyR-based systems it is expected to support stable gain, synchrony, and fidelity of Ca^2+^ release. This links the conserved molecular geometry of the activation site to the reliability of Ca^2+^ signaling across diverse physiological settings. This geometry-set quantum noise feeding a regenerative amplifier could be a general principle of biology spanning multiple fields from inner ear biophysics, vertebrate segmentation, plant guard cell signaling, comparative cardiac physiology, and stochastic thermodynamics of ion channels.

## Methods

### Tunneling pipeline

A four-step pipeline estimates the per-channel noise amplitude from quantum mechanics and cryo-EM structural data (SI Appendix, Methods). (1) Donor-acceptor geometry was characterized across 12 cryo-EM structures; two pK_*a*_ predictors bracket E3922 protonation at 3–65% at pH 7.0. (2) The tunneling splitting *ΔE* was computed from the 1D Schrödinger equation for a proton in a quartic double-well:

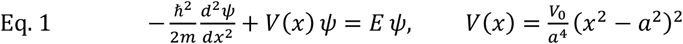

(3) Thermal O⋯O fluctuations (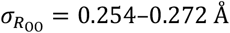, mean 0.264 Å across six open structures; bounded below by Herschlag limit ≥0.06 Å) were modeled as a two-tier Ornstein-Uhlenbeck process and converted to noise via κ(*t*) = tanh[(*ΔE*(*t*) − *μ*)/(2*μ*)]. (4) After coarse-graining (*τ*_slow_ = 1 ms, *f*_slow_ = 0.4, *Δ* = 4 ms), *σ*_κ,eff_ ≈ 0.20.

### Pacemaker cell model

Each cell is described by the Maltsev–Lakatta rabbit SAN model [33], tracking 29 coupled state variables. The membrane potential is governed by:

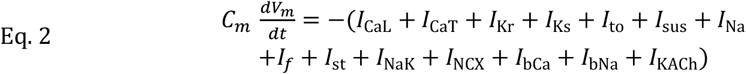

where *Cm* = 20 pF. Quantum noise enters as a per-CRU multiplicative modulation:

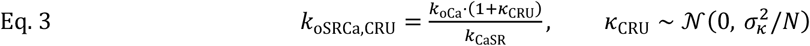

updated every 4 ms. Monte Carlo spark ignition is present in all conditions including the control (*κ* = 0). CRU sizes were drawn from the super-resolution imaging distribution [32] with RyR density ρ = 67 RyR/µm^2^. Temperature-dependent rates were scaled by 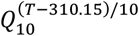 (gating: 2.5; SERCA: 2.6; diffusion: 1.3; NCX: 2.0; NaK: 2.0; spark inactivation: 2.7; SI Appendix, Methods). The RyR2 opening rate *k*_oCa_ was *not* scaled (*Q*_10_ = 1.0; [20]). Three conditions were compared: (1) quantum (*σ*_*κ*_ = 0.20, constant); (2) classical (*σ*_*κ*_(*T*) = 0.20 × 2.7^(*T*−310.15)/10^); (3) control (*κ* = 0).

### Experimental Ca^2+^ recordings

Single spontaneously beating SAN cells were isolated from rabbit hearts [41] and loaded with the Ca^2+^ indicator Fluo-4. Whole-cell Ca^2+^ fluorescence was recorded at 100 frames/s using a high-speed CCD camera. Each cell (*n* = 17) was recorded at both 25 ^∘^C and 37 ^∘^C in a paired design. The CV of inter-transient intervals and mean beat rate were compared using the paired t-test (SI Appendix, Methods).

## Data Availability Statement

Analysis scripts, tunneling pipeline code, and notebooks for reproducing all computational results are available at https://github.com/victoramaltsev/RyR-network-SANC-model. Cryo-EM structures used in this study are publicly available from the RCSB Protein Data Bank (https://www.rcsb.org/) under accession codes 7UA9, 7UA5, 7U9R, 7U9Q, 5GOA, 6JG3, 6JGZ, 6JIY, 6JH6, 6JHN, 7VML, and 5TAL.

## Author Contributions

AM designed the study, developed the tunneling pipeline and single-cell noise implementation, performed the computations, and wrote the manuscript. EGL contributed to the conceptual framework and physiological interpretation. VAM supervised the project, performed experiments, and contributed to the theoretical framework.

## Competing Interests

The authors declare no competing interests.

## Funding

This research was supported by the Intramural Research Program of the National Institutes of Health (NIH). The contributions of the NIH authors are considered Works of the United States Government. The findings and conclusions presented in this paper are those of the authors and do not necessarily reflect the views of the NIH or the U.S. Department of Health and Human Services.

## Ethics Statement

The animal study conformed to the Guide for the Care and Use of Laboratory Animals, published by the US National Institutes of Health. The experimental protocols were approved by the Animal Care and Use Committee of the National Institute on Aging, National Institutes of Health.

## Acknowledgments

The authors thank Bruce D. Ziman for skillful isolation of single sinoatrial node cells and Larissa A. Maltseva for skilled assistance in the Ca^2+^ imaging experiments.

## SI Appendix

### SI Appendix: Methods

#### E3922 protonation establishes a donor-acceptor pair

PROPKA pK_*a*_ prediction [1] was applied to 12 deposited structures spanning human and porcine RyR2 in open/closed states plus rabbit RyR1 as a cross-isoform check (Table S4). In each case, one complete protomer plus all inter-chain residues within 20 Å of the Ca^2+^ activation site were retained, and Ca^2+^ ions were included where modeled. PROPKA treats metal ions as point charges in its empirical electrostatic model. Across all 12 structures, the E3922-equivalent pK_*a*_ ranges from 5.15 to 10.19, with 9 of 12 structures showing pK_*a*_ > 7 (majority protonated at pH 7.0). The dephosphorylated open state (7UA9) gives pK_*a*_ = 7.27. However, pKAI+ predicts the reverse for 7UA9 (E3922 = 5.55, E3848 = 6.35; Table S4), so the cross-method support for the specific donor assignment depicted in Fig. 4A is weaker than either predictor alone would suggest. In the rabbit RyR1 structure (5TAL), the donor-acceptor roles are swapped. E3893 (the E3848 equivalent) has pK_*a*_ = 8.46, while E3967 (E3922 equivalent) has pK_*a*_ = 5.47. The O⋯O geometry is similar, but the residue with the higher pK_*a*_ (and thus the likely proton donor) differs.

As a cross-method check, the deep-learning predictor pKAI+ [2], trained on Poisson-Boltzmann calculations rather than empirical rules, was applied to the same structures. pKAI+ reads protein ATOM records without modeling metal ions. For 7UA9 it predicts E3922 pK_*a*_ = 5.52, elevated 1.3 units above the intrinsic glutamate value (4.20) by burial alone. Both methods agree that burial elevates E3922 pK_*a*_ above intrinsic, and differ in magnitude because PROPKA additionally models Ca^2+^ as a point charge. The two methods bracket plausible values: pKAI+ at the lower end (∼ 5.5, ∼ 3% protonated) and PROPKA at the upper end (∼ 7.3, ∼ 65% protonated). Experimental measurements in the calbindin D_9k_ Ca^2+^ binding site [3], where glutamate pK_*a*_ values reach ∼ 6.5 under comparable burial, fall between the two estimates.

Near-physiological pK_*a*_ values for ion channel carboxylates have independent precedent. Free-energy calculations for the NavMs selectivity filter found glutamate pK_*a*_ values of 6.7– 7.5 [4]. Even at the pKAI+ lower end (∼ 3% protonated), a donor-acceptor pair exists a fraction of the time sufficient for transient proton transfer. Constant-pH molecular dynamics or neutron crystallography at the activation site would further constrain the protonation state.

Across open-state structures, the shortest O⋯O distances, 2.74 Å (E–T, 5GOA) and 2.88 Å (E–E, 6JIY), approach the range where neutron crystallography has confirmed proton sharing (*R*_OO_ = 2.43–2.55 Å; [5,6,7,8]). These distances require only 0.19–0.33 Å of thermal compression, corresponding to 0.7–1.3*σ* events sampled 10–24% of the time at the Boltzmann rotamer-derived 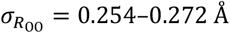 (mean 0.264 Å across six open structures; bounded below by Herschlag limit ≥0.06 Å). This follows the “tunneling-ready state” approach from enzyme catalysis [9]. Three additional pathways could supplement direct E3922–E3848 transfer: (1) water-mediated proton relay through the hydrated activation site; (2) tunneling along the shorter E3848–T4931 coordinate (2.99 Å); and (3) tunneling during the Ca^2+^ binding transition, when carboxylate geometry may transiently compress. Each pathway is consistent with the pipeline’s quantitative predictions, because *σ*_*κ*_ depends on the transfer function and OU parameters rather than on the specific donor-acceptor pair. Site-specific QM/MM calculations will be needed to determine which pathway is operative.

#### Tunneling splitting, thermal fluctuations, and the saturating transfer function

Equation 1 (main text) was solved numerically using a finite-difference method on *N* = 2,500 grid points spanning *x* ∈ [−3*a*, +3*a*], yielding a tridiagonal Hamiltonian matrix. The lowest six eigenvalues were obtained via the Lanczos algorithm. The tunneling splitting *ΔE* = *E*_1_ − *E*_0_ is exponentially sensitive to barrier width (∼ 2.9 decades per Å). At the reference parameters (*V*_0_ = 20 kJ/mol, *R*_OO_ = 2.60 Å), *ΔE*_H_ = 103.3 cm^−1^. At the cryo-EM distance (*R*_OO_ = 2.99 Å), *ΔE*_H_ = 7.2 cm^−1^. The solver was validated against the formic acid dimer benchmark.

Thermal fluctuations in *R*_OO_ were modeled as an Ornstein-Uhlenbeck process:

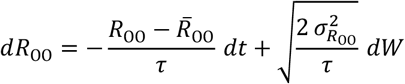

with 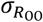 bounded by: (i) ANM analysis (∼ 0.007 Å, backbone only); (ii) Ca^2+^ -coordination-weighted Boltzmann rotamer sampling of the E3848–E3922 sidechain pair across all six open H-bonding structures (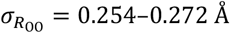, mean 0.264 ± 0.006 Å; 500,000 configurations per structure weighted by coordination energy (*k* = 50 kJ/mol/Å^2^), screened Coulomb interactions (*ϵ*_*r*_ = 10), and steric repulsion at *T* = 310 K, with H-bond filter *d* < 3.5 Å, convergent across three independent chi-angle width scenarios); (iii) Herschlag thermodynamic lower bound (≥ 0.06 Å; [10]); (iv) crystallographic survey [11]. The consistency of 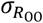 across structures (SD = 0.006 Å) indicates that the rotameric freedom is determined by glutamate sidechain geometry and Ca^2+^ coordination constraints rather than by the equilibrium O⋯O distance.

Protein dynamics contribute on two timescales: fast breathing modes (*τ*_fast_ ∼ 10 *μ*s) and slow conformational substate transitions (*τ*_slow_ ∼ 0.1–5 ms; [12,13,14]). This is modeled as a two-tier process:

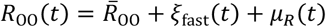

where 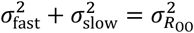 and 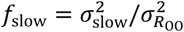. A structural survey of 12 cryo-EM structures provides site-specific evidence for slow conformational dynamics at this locus (*f*_slow_ > 0). Among 8 open-state structures, the E3848–E3922 O⋯O distance splits into two discrete macrostates: six structures in H-bonding range (2.88–3.82 Å, mean 3.31 ± 0.35 Å) and three in a non-H-bonding state (4.58–6.19 Å). B-factor analysis shows the site is well-ordered within each structure (*B*_site_/*B*_global_ = 0.57 ± 0.16; *p* < 10^−3^; Table S3).

Ca^2+^-coordination-weighted Boltzmann rotamer sampling across all six open H-bonding structures yields 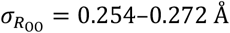 (mean 0.264 ± 0.006 Å). The saturating transfer function (tanh) ensures bounded output, consistent with the finite dynamic range of ion channel gating. Alternative monotone mappings (linear, logarithmic) yield higher amplitudes that also fall within the coherence resonance window. The saturating form gives the lowest value among the three. After two-tier coarse-graining (*τ*_slow_ = 1 ms, *f*_slow_ = 0.4), the full nonlinear pipeline at the structural mean 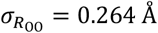 yields *σ*_κ,eff_ ≈ 0.20, approximately temperature-invariant (Figure 4, main text). The value *f*_slow_ = 0.4 reflects the physical constraint imposed by Ca^2+^ coordination at the activation site. The divalent cation bridges both carboxylates (E3848 and E3922), creating a strong harmonic stiffness penalty (∼ 50–100 kJ/mol/Å^2^) for any rearrangement that disrupts the coordination geometry. Fast local vibrations (bond stretching, angle bending) are small enough to remain within the coordination well, but slow conformational rearrangements (sidechain rotamer transitions, backbone shifts) must overcome this penalty and are therefore suppressed relative to an unconstrained protein site. The B-factor ratio *B*_site_/*B*_global_ = 0.57 confirms that the activation site is more ordered than the protein average, consistent with Ca^2+^-mediated rigidity that preferentially suppresses the slow tier. The cryo-EM macrostates (H-bonding vs. non-H-bonding, Table S3) correspond to Ca^2+^-bound vs. Ca^2+^-free gating transitions, which are distinct from the thermal fluctuations within the Ca^2+^-bound open state modeled here. The dose-response scan (Table S6) shows that CV_IBI_ reaches a minimum at *σ*_*κ*_ ≈ 0.20–0.22. A sensitivity analysis across the plausible parameter space (Table S6) shows that *σ*_κ,eff_ falls within this window for moderate values of *τ*_slow_ and *f*_slow_, but exceeds it when both parameters are large (*τ*_slow_ ≥ 5 ms and *f*_slow_ ≥ 0.7). This constraint is consistent with the Ca^2+^ coordination argument: the high-*f*_slow_ regime, in which most distance variance comes from slow conformational rearrangements, is physically disfavored at a Ca^2+^-stabilized site. Site-specific QM/MM calculations at the RyR2 activation site will further constrain *τ*_slow_ and *f*_slow_. Uncertainties across pipeline steps do not multiply independently because the saturating transfer function compresses large variations in tunneling splitting into bounded variations in *σ*_*κ*_: a 10× range in *R*_OO_-dependent splitting produces <1.3× variation in *σ*_*κ*_.

#### Two-tier conformational dynamics preserve noise through the 4 ms update window

The simulation updates κ every *Δ* = 4 ms, so the relevant quantity is the block-averaged modulation 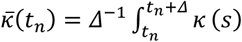 *ds*. For a single-tier OU with *τ*_fast_ = 10 *μ*s, variance attenuates by 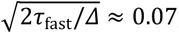, reducing *σ*_*κ*_ to ∼ 0.023. In the two-tier model, the slow tier (*τ*_slow_ ∼ 1–5 ms, comparable to *Δ*) is largely preserved. At the structural mean 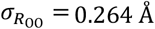 with *τ*_slow_ = 1 ms and *f*_slow_ = 0.4, the full nonlinear pipeline yields *σ*_κ,eff_ ≈ 0.20, approximately temperature-invariant (Figure 4, main text). This temperature invariance arises because the saturating transfer function absorbs the modest 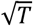 scaling of 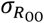. Thermal fluctuations in the donor-acceptor distance scale as 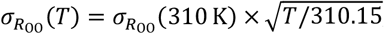, giving 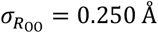 at 5 C (278 K) and 0.266 Å at 42 C (315 K), a variation of only ±3% from the 37 ^∘^C reference. Because the tunneling splitting depends exponentially on *R*_OO_ but the transfer function saturates, this input variation produces <1% variation in *σ*_κ,eff_ across the full 5–42 ^∘^C range, confirming that the effective noise amplitude is essentially temperature-invariant.

The dose-response scan (Table S6) shows that this amplitude range produces coherence resonance in the single-cell model. The simulation draws independent Gaussian samples at each 4 ms step, which does not preserve temporal correlations between successive windows. For the slow tier with *τ*_slow_ ∼ 1–5 ms, adjacent block averages of the true OU process would retain autocorrelation *ρ*_adj_ ≈ 0.2–0.5. Coherence resonance is a generic property of noise-driven excitable systems whose existence is robust to noise color [15]. Explicit OU-filtered simulations will characterize the quantitative shift, if any, in the resonance optimum. The per-channel modulation *k*_oCa_ indirectly through fluctuations in local [Ca^2+^]_sub_, and its amplitude depends on the underlying transition rates. If *k*_oCa_ is temperature-independent, the stochastic component it generates is also temperature-independent; however, the quantum-tunneling mechanism provides a physical basis for why the opening rate carries this property.

#### Supercluster ignitions set the cell-level noise amplitude

Each CRU’s noise amplitude is 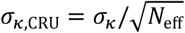, where *N*_eff_ = *N*/(1 + (*N* − 1)*ρ*) and *ρ* is the inter-channel correlation. The cell-level noise involves two stages:

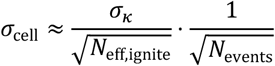

In the independent-channel limit (*ρ* = 0) at *σ*_κ,eff_ = 0.20, *σ*_cell_ ≈ 0.004. In the strong-correlation limit (*ρ* → 1, each ignition carries full per-channel noise), 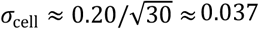. With fewer dominant events (*N*_events_ ≈ 16), the upper bound extends to ∼ 0.050.

#### Proton-Ca^2+^ exchange controls gating in SR membrane proteins

RyR2 is directly pH-sensitive, with half-inhibition at pH ∼ 6.5 on the cytoplasmic side [16,17]. This is only 0.5–1.0 pH units above physiological pH, making the channel highly sensitive to local proton concentration changes. [18] demonstrated that SERCA countertransports 2–3 H^+^ per Ca^2+^ cycle through protonation of coordinating carboxylates. This establishes that proton–Ca^2+^ exchange directly controls conformational gating in SR membrane proteins. DFT calculations show that protonating even one carboxylate in a Ca^2+^ binding site reduces binding affinity by potentially >100-fold [19,20]. NMDA receptor proton inhibition requires only ∼ 2.7 kcal/mol to shift the gating equilibrium 100-fold [21]. A neutron crystallography study of a different Ca^2+^-carboxylate binding site found all coordinating carboxylates deprotonated, but also revealed a low-barrier hydrogen bond (LBHB) between fucose hydroxyl and Asp96 induced by two Ca^2+^ ions [22]. Site-specific calculations of how Ca^2+^ modifies the proton transfer barrier will further refine the pipeline. As an analogy, a neighboring acidic residue can cooperatively strengthen a short strong hydrogen bond in ATP-synthase/bedaquiline binding [23], illustrating how local electrostatics can reshape proton-sharing energetics at carboxylate sites.

#### Temperature scaling of the single-cell model

The Maltsev–Lakatta single-cell SAN model [24,25] was originally parameterized at 37 ^∘^C (*T* = 310.15 K). To simulate other temperatures, all rate-dependent processes except the RyR2 opening rate were scaled using a *Q*_10_ formulation:

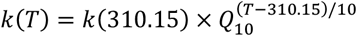

Six groups of processes were assigned distinct *Q*_10_ values based on experimental data (Table S1).

The RyR2 opening rate *k*_oCa_ was not scaled (*Q*_10_ = 1.0), based on the near-zero activation energy measured by [26]. The Nernst potentials were recalculated at each temperature via *RT*/*F*. Gating steady-state curves (voltage-dependent *α* and *β* rate constants) were not shifted along the voltage axis; only the time constants were scaled. This is a standard approximation that assumes the voltage dependence of gating reflects the electric field across the membrane, which does not change with temperature, while the kinetic rates reflect thermal activation, which does.

Quantum noise was implemented as a per-CRU multiplicative modulation of *k*_oSRCa_ (Eq. 3, main text), updated every 4 ms (∼ 533 integration ticks at *Δt* = 0.0075 ms). At each update, each CRU drew a new κ_CRU_ from 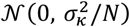, where *N* is the number of RyR channels in that CRU and *σ*_*κ*_ = 0.20. The 4 ms update interval corresponds to the timescale of Ca^2+^ spark rise and submembrane Ca^2+^ integration, below which the cell cannot respond to individual tunneling events.

Three noise conditions were compared at each temperature: (1) quantum (*σ*_*κ*_ = 0.20, constant across all temperatures); (2) classical (*σ*_*κ*_(*T*) = 0.20 × 2.7^(*T*−310.15)/10^, declining with cooling); (3) control (*κ* = 0, no additional noise beyond the intrinsic Monte Carlo stochasticity of spark ignition, which is present in all conditions). Simulations ran for 30,000 ms at 25 ^∘^C and 30,000 ms at 37 ^∘^C. The first 5,000 ms were discarded as warmup in all cases.

#### Isolated SAN cells maintain Ca^2+^ transients at 25 and 37 ^∘^C

Single, spontaneously beating, spindle-shaped SAN cells were isolated from 8–12 week old male New Zealand White rabbit hearts as previously described [27] in accordance with NIH guidelines for the care and use of animals, protocol # 457-LCS-2024. Intracellular Ca^2+^ dynamics were measured by 2D imaging of fluorescence emitted by the Ca^2+^ indicator Fluo-4 using a high-speed Hamamatsu C9100-12 CCD camera (100 frames/s, 512 × 512 pixels, 8.192 mm square sensor) mounted on a Zeiss Axiovert 100 inverted microscope with a ×63 oil immersion lens and a fluorescence excitation light source (Sutter Instrument, LB-LS/Q17) housing a 175W xenon lamp. Fluo-4 fluorescence excitation (blue light, 470/40 nm) and emission light collection (green light, 525/50 nm) were performed using the Zeiss filter set 38 HE. Cells were loaded with 1.5 *μ*M Fluo-4AM (Sigma-Aldrich) for 10 min at room temperature. Fluo-4AM was subsequently washed out of the chamber, and Ca^2+^ signals were measured within the ensuing 60 min at 25 °C / 37 ^∘^C ± 0.1 ^∘^C.

The physiological bathing solution contained (in mM): NaCl 140, KCl 5.4, MgCl_2_ 2, HEPES 5, CaCl_2_ 1.8, pH 7.3 (adjusted with NaOH). Temperature was controlled by an Analog TC2BIP 2/3Ch bipolar temperature controller (CellMicroControls, USA), heating both the glass bottom of the perfusion chamber and the solution entering the chamber via a pre-heater. Each cell (*n* = 17) was recorded at both 25 ^∘^C and 37 ^∘^C in a paired design, with order counterbalanced across cells.

Ca^2+^ signal waveforms were generated from movies (stacks of consecutive images) of Ca^2+^-dependent fluorescence of Fluo-4 by calculating the spatial average fluorescence of each image within a user-defined region of interest (ROI) encompassing the entire cell perimeter, yielding a whole-cell Ca^2+^ signal time series at 10 ms resolution. Ca^2+^ transient peaks were detected using prominence-based peak finding (SciPy `find_peaks`) after removing the initial fluorescence plateau via rolling-standard-deviation onset detection. The CV of inter-transient intervals and mean beat rate were computed for each cell at each temperature. Statistical comparison used the paired t-test.

**Table S1:**
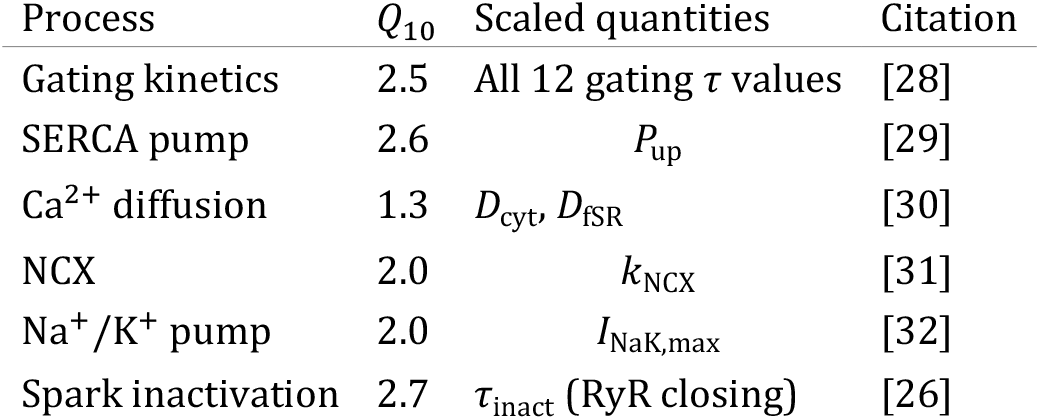
Q_10_ scaling applied to biophysical processes in the temperature sweep simulations.

**Table S2:** Glossary of symbols and abbreviations.

| Symbol | Definition |
| --- | --- |
| $C_m$ | Membrane capacitance (20 pF) |
| CICR | Ca<sup>2+</sup>-induced Ca<sup>2+</sup> release |
| CRU | Ca<sup>2+</sup> release unit (cluster of RyR2 channels) |
| CV<sub>IBI</sub> | Coefficient of variation of inter-transient intervals (inter-beat intervals in simulations) |
| $\Delta E$ | Tunneling splitting (energy gap between ground and first excited proton states) |
| ICC | Interstitial cells of Cajal (gut pacemaker cells) |
| $I_{\text{KACH}}$ | Acetylcholine-sensitive potassium current |
| $k_{\text{oCa}}$ | Ca<sup>2+</sup>-dependent RyR2 opening rate |
| $k_{\text{iCa}}$ | Ca<sup>2+</sup>-dependent RyR2 inactivation rate |
| $k_{\text{oSRCa}}, k_{\text{iSRCa}}$ | SR-load-modulated opening and inactivation rates |
| $\kappa_i$ | Per-CRU quantum noise modulation factor, time-dependent |
| $N$ | Number of RyR2 channels per CRU |
| NCX | Na<sup>+</sup>/Ca<sup>2+</sup> exchanger |
| $P_{\text{up}}$ | SERCA uptake rate (mM/s) |
| $R_{\text{OO}}$ | Donor-acceptor O...O distance (Å) |
| $\rho$ | RyR density (RyR/$\mu\text{m}^2$) |
| RyR1, RyR3 | Skeletal and brain ryanodine receptor isoforms |
| RyR2 | Cardiac ryanodine receptor |
| SAN | Sinoatrial node |
| $\sigma_\kappa$ | Per-channel noise amplitude (dimensionless) |
| $\sigma_{\text{cell}}$ | Per-cell noise amplitude |
| $\sigma_{R_{\text{OO}}}$ | RMS amplitude of donor-acceptor distance fluctuations (Å) |
| SR | Sarcoplasmic reticulum |
| $V_0$ | Proton transfer barrier height (kJ/mol) |

**Table S3:** Structural survey of O⋯O distances at the RyR Ca^2+^ activation site across 12 cryo-EM structures. B_site_/B_global_: ratio of activation site B-factor to protein-wide mean; values < 1 indicate the site is more ordered than average.

| PDB | Species | State | Res. (Å) | E-E (Å) | $\bar{B}$ | $\sigma_{pos}$ (Å) | H-bond? | $B_{site}/B_{global}$ |
| --- | --- | --- | --- | --- | --- | --- | --- | --- |
| 7UA9 | Human | Open | 3.6 | 3.12 | 77.5 | 0.99 | Yes | 0.62 |
| 5GOA | Porcine | Open | 4.2 | 3.82 | 78.3 | 0.99 | Yes | 0.39 |
| 6JG3 | Porcine | Open | 4.0 | 3.45 | 138.5 | 1.33 | Yes | 0.47 |
| 6JGZ | Porcine | Open | 4.0 | 3.26 | 132.4 | 1.30 | Yes | 0.52 |
| 6JIY | Porcine | Open | 3.9 | 2.88 | 56.3 | 0.84 | Yes | 0.46 |
| 6JH6 | Porcine | Open | 4.8 | 4.58 | 152.7 | 1.39 | No | 0.55 |
| 6JHN | Porcine | Open | 4.0 | 5.77 | 188.1 | 1.54 | No | 0.60 |
| 7U9R | Human | Open | 3.7 | 6.19 | 112.2 | 1.19 | No | 1.01 |
| 7UA5 | Human | Closed | 2.8 | 5.36 | 53.6 | 0.82 | No | 0.53 |
| 7U9Q | Human | Closed | 3.1 | 6.02 | 74.4 | 0.97 | No | 0.62 |
| 7VML | Human | Closed | 3.0 | 5.13 | 33.3 | 0.65 | No | 0.51 |
| 5TAL | Rabbit RyR1 | Open | 3.8 | 3.01 | 84.5 | 1.03 | Yes | 0.51 |

**Table S4:** PROPKA and pKAI+ pK_a_ predictions for the E3922-equivalent and E3848-equivalent glutamates across 12 cryo-EM structures spanning three species and both RyR isoforms. PROPKA treats metal ions as point charges; pKAI+ reads only protein ATOM records and does not model metal ions. Porcine residue numbering: +1 offset (E3849/E3923). RyR1 (5TAL): E3967/E3893. % prot: PROPKA-based protonation at pH 7.0.

| PDB | Species/State | Ca<sup>2+</sup> | Res. (Å) | PROPKA | pKAI+ | E3922 | E3848 | %<br>prot |
| --- | --- | --- | --- | --- | --- | --- | --- | --- |
|  |  |  |  | E3922 | E3848 |  |  |  |
| *Human RyR2* |  |  |  |  |  |  |  |  |
| 7UA9 | Open, dephos | Yes | 3.59 | 7.27 | 4.40 | 5.55 | 6.35 | 65 |
| 7UA5 | Closed, dephos | No | 2.83 | 7.72 | 5.30 | 5.24 | 3.27 | 84 |
| 7U9R | Open, PKA | Yes | 3.69 | 5.15 | 5.30 | 6.36 | 4.04 | 1 |
| 7U9Q | Closed, PKA | No* | 3.11 | 8.17 | 5.56 | 4.53 | 4.26 | 94 |
| *Porcine RyR2* |  |  |  |  |  |  |  |  |
| 5GOA | Open | Yes | 4.20 | 10.19 | 6.98 | 5.55 | 5.42 | 100 |
| 6JG3 | Open | Yes | 4.00 | 9.31 | 6.26 | 3.22 | 4.90 | 100 |
| 6JGZ | Open | Yes | 4.00 | 9.65 | 6.31 | 6.61 | 4.90 | 100 |
| 6JIY | Open | Yes | 3.90 | 8.31 | 4.92 | 6.04 | 5.12 | 95 |
| 6JH6 | Open | Yes | 4.80 | 8.46 | 5.95 | 6.03 | 4.68 | 97 |
| 6JHN | Open | Yes | 4.00 | 5.90 | 6.59 | 5.88 | 4.97 | 7 |
| *Human RyR2 (closed, Bhatt et al.)* |  |  |  |  |  |  |  |  |
| 7VML | Closed | No | 3.00 | 7.60 | 5.55 | 4.74 | 3.96 | 80 |
| *Rabbit RyR1 (cross-isoform)* |  |  |  |  |  |  |  |  |
| 5TAL | Open | Yes | 3.80 | 5.47 | 8.46 | 6.47 | 6.15 | 3 |

\*7U9Q is annotated as open in the PDB deposition but lacks modeled Ca<sup>2+</sup> at the activation site, suggesting incomplete Ca<sup>2+</sup> occupancy or a transitional state.

**Table S5:** PROPKA pK_a_ decomposition for E3922 in 7UA9 (open, dephosphorylated, Ca^2+^ modeled).

| Contribution | $pK_a$<br>shift | Notes |
| --- | --- | --- |
| Model (solution) $pK_a$ | 4.50 | PROPKA3 reference for Glu |
| Desolvation / burial | +2.65 | 94% side-chain burial |
| Hydrogen bonds | +1.56 | Primarily E3848 side chain |
| Coulombic interactions | -1.44 | $Ca^{2+}$ (-2.92), offset by E3848 (+1.22), E4932 (+0.41) |
| **Predicted $pK_a$** | **7.27** |  |

**Table S6:** Coherence resonance dose-response at 37 ^∘^C. Noise amplitude *σ*_*κ*_ was varied across nine conditions. CV_IBI_ reaches a minimum at *σ*_*κ*_ = 0.20 and rises at both lower and higher amplitudes, tracing the U-shaped signature of coherence resonance. All runs used the same CRU realization (realistic CRU distribution [25]); first 5 s discarded as warmup. The three-condition temperature comparison (Results) used matched parameters but independent 30 s runs at each temperature.

| Condition | $\sigma_\kappa$ | $CV_{\text{IBI}}$ |
| --- | --- | --- |
| Control ($\kappa = 0$) | — | 0.028 |
| $\sigma_\kappa = 0.12$ | 0.12 | 0.028 |
| $\sigma_\kappa = 0.14$ | 0.14 | 0.028 |
| $\sigma_\kappa = 0.16$ | 0.16 | 0.029 |
| $\sigma_\kappa = 0.18$ | 0.18 | 0.030 |
| $\sigma_\kappa = 0.20$ | 0.20 | **0.026** |
| $\sigma_\kappa = 0.22$ | 0.22 | 0.026 |
| $\sigma_\kappa = 0.24$ | 0.24 | 0.029 |
| $\sigma_\kappa = 0.26$ | 0.26 | 0.031 |

## Notes

https://github.com/victoramaltsev/RyR-network-SANC-model

### Competing Interest Statement

The authors have declared no competing interest.

